# The *Enterprise*: A massive transposon carrying *Spok* meiotic drive genes

**DOI:** 10.1101/2020.03.25.007153

**Authors:** Aaron A. Vogan, S. Lorena Ament-Velásquez, Eric Bastiaans, Ola Wallerman, Sven J. Saupe, Alexander Suh, Hanna Johannesson

## Abstract

The genomes of eukaryotes are full of parasitic sequences known as transposable elements (TEs). Most TEs studied to date are relatively small (50 – 12000 bp), but can contribute to very large proportions of genomes. Here we report the discovery of a giant tyrosine-recombinase-mobilized DNA transposon, *Enterprise*, from the model fungus *Podospora anserina*. Previously, we described a large genomic feature called the *Spok* block which is notable due to the presence of meiotic drive genes of the *Spok* gene family. The *Spok* block ranges from 110 kb to 247 kb and can be present in at least four different genomic locations within *P. anserina*, despite what is an otherwise highly conserved genome structure. We have determined that the reason for its varying positions is that the *Spok* block is not only capable of meiotic drive, but is also capable of transposition. More precisely, the *Spok* block represents a unique case where the *Enterprise* has captured the *Spok*s, thereby parasitizing a resident genomic parasite to become a genomic hyperparasite. Furthermore, we demonstrate that *Enterprise* (without the *Spoks*) is found in other fungal lineages, where it can be as large as 70 kb. Lastly, we provide experimental evidence that the *Spok* block is deleterious, with detrimental effects on spore production in strains which carry it. In contrast to the selfish role of the *Enterprise* in *P. anserina*, we hypothesize that the mobility of the *Enterprise* may also play an adaptive role in fungi when *Enterprise* undergoes horizontal transfer while carrying metabolic genes. This union of meiotic drivers and a transposon has created a selfish element of impressive size in *Podospora*, challenging our perception of how TEs influence genome evolution and broadening the horizons in terms of what the upper limit of transposition may be.

## Introduction

Transposable elements (TEs) are major agents of change in eukaryotic genomes. Their ability to selfishly parasitise their host replication machinery has large impacts on both genome size and on gene regulation (Chénais et al. 2012). In extreme cases, TEs can contribute up to 85% of genomic content (Schnable et al. 2009) and expansion and reduction of TEs can result in rapid changes in both genome size and architecture (Haas et al. 2009; Talla et al. 2017; Möller and Stukenbrock 2017). Generally, TEs have small sizes (∼50 – 12000 bp) and accomplish these large-scale changes through their sheer number. For example, there are ∼1.1 million *Alu* elements in the human genome, which have had a large impact on genome evolution (Bennett et al. 2008; Jurka 2004). The largest known cases among Class I retrotransposons are long terminal repeat (LTR) elements that can be as large as 30 kb, but among Class II DNA transposons, Mavericks/Polintons are known to grow as large as 40 kb through the capture of additional open reading frames (ORFs) (Arkhipova and Yushenova 2019). Recently, a behemoth TE named *Teratorn* was described in teleost fish; it can be up to 182 kb in length, dwarfing all other known TEs. *Teratorn* has achieved this impressive size by fusing a *piggyBac* DNA transposon with a herpesvirus, thereby blurring the line between TEs and viruses (Inoue et al. 2017, 2018). Truly massive transposons may be lurking in the depths of many eukaryotic genomes, but the limitations of short-read genome sequencing technologies and the lack of population-level high-quality assemblies may make them difficult to identify.

The *Spok* block is a large genomic feature that was first identified thanks to the presence of the s**po**re **k**illing (*Spok*) genes in species from the genus *Podospora* (Vogan, Ament-Velásquez, et al. 2019; Grognet et al. 2014). The *Spoks* are selfish genetic elements that bias their transmission to the next generation in a process known as meiotic drive. Here, drive occurs by inducing the death of spores that do not inherit them, through a single protein that operates as both a toxin and an antidote (Vogan, Ament-Velásquez, et al. 2019; Grognet et al. 2014). The first *Spok* gene described, *Spok1*, was discovered in *P. comata* (Grognet et al. 2014). In *P. anserina*, the homologous gene *Spok2* is found at high population frequencies, while two other genes of the *Spok* family, *Spok3* and *Spok4*, are at low to intermediate frequencies (Vogan, Ament-Velásquez, et al. 2019). Unlike *Spok1* and *Spok2*, however, *Spok3* and *Spok4* are always associated with a large genomic region (the *Spok* block). The *Spok* block can be located at different chromosomal locations in different individuals, but is never found more than once in natural strains. The number of *Spok* genes and the location of the *Spok* block (which carries *Spok3, Spok4* or both) define the overall meiotic driver behavior of a given genome, which can be classified into the so-called ***P****odospora* **s**pore **k**illers or *Psk*s (Vogan, Ament-Velásquez, et al. 2019; van der Gaag et al. 2000). The *Spok* block stands out not only because of its size, typically around 150 kb, but also because there is otherwise high genome synteny among strains of *P. anserina* and with the related species *P. comata* and *P. pauciseta* (Vogan, Ament-Velásquez, et al. 2019).

The fact that the *Spok* block is found at unique genomic positions between otherwise highly similar strains is of prime interest as each novel *Spok* block position creates a unique meiotic drive type (*Psk*) due to the intricacies of meiosis in *Podospora*. In this study, we have determined that the reason the *Spok* block occurs at multiple genomic locations is that the *Spok* block itself is actually a unique version of the *Enterprise*, a novel DNA transposon that is mobilized by a tyrosine recombinase (YR). The *Spok* block represents an *Enterprise* that has captured meiotic drive genes and subsequently grown to a massive size through a gradual accumulation of DNA sequence. We find copies of *Enterprise* without meiotic drive genes in other fungal species that are at least 70 kb, placing *Enterprise* among the largest known TEs, even when not harbouring the *Spoks*. The *Spok* block version of *Enterprise* not only represents the largest TE discovered to date (reaching 247 kb), but is also capable of both transposition and meiotic drive. Furthermore, our data suggest that the *Spok* block is associated with fitness costs in *P. anserina*. This discovery drastically changes what we know about the size limits of transposition, and provides new insight into the role of massive TEs in eukaryotic genome evolution.

## Results

### Newly isolated *Podospora* strains reveal a novel spore killer type

We isolated and sequenced with Illumina HiSeq technology a strain of *P. anserina* (named Wa137) and two strains of *P. comata* (Wa131 and Wa139) collected in Wageningen, the Netherlands. The strains Wa137 and Wa139 were also sequenced using MinION Oxford Nanopore technology, achieving similar quality to published chromosome-level assemblies (**Supplementary Table 1**). Additionally, we included in our analyses all published long-read genomes of *P. anserina* (10 strains), *P. pauciseta* (one), and *P. comata* (one) (Espagne et al. 2008; Vogan, Ament-Velásquez, et al. 2019; Silar et al. 2019), which are mostly assembled at chromosome level, with respective genome sizes around 36 Mb (**Supplementary Table 1**). Previous work demonstrated that the *Spok* block can be found at three unique chromosomal positions among characterized *P. anserina* strains, defining the killer types *Psk-1*/*5* (with the *Spok* block located at chromosome 3), *Psk-2* (right arm of chromosome 5) and *Psk-7/8* (left arm of chromosome 5) (Vogan, Ament-Velásquez, et al. 2019) (**Table 1**). The newly isolated strain Wa137 was found to have a *Spok* block with the largest size yet reported (247 kb) and in a novel position (chromosome 1), conferring it a new spore killing phenotype that we named *Psk-9* (see Vogan, Ament-Velásquez, et al. (2019) for the principle of nomenclature). The published *P. pauciseta* genome has a *Spok* block in chromosome 4 (*Psk-P1*), but it seems to represent a fragmented version of the *P. anserina* blocks (Vogan, Ament-Velásquez, et al. 2019). Unlike *P. anserina* and *P. pauciseta*, the newly isolated strains of *P. comata* were found to have a single full-copy *Spok* gene (*Spok1*) which is not associated with any *Spok* block-like features, in agreement with the previously published reference genome of *P. comata* (**Table 1**) (Grognet et al. 2014; Silar et al. 2019).

**Table 1.** List of strains used in this study.

| Strain | Species | *Psk*<sup>a</sup> | Genomic location | Size | *Enterprise*<sup>b</sup> | *Spok* genes |
| --- | --- | --- | --- | --- | --- | --- |
| Wa21 | *P. anserina* | 2 | Chr5 R | 142433 | *Spok* block | *Spok2 Spok3* |
| Wa28 | *P. anserina* | 2 | Chr5 R | 161297 | *Spok* block | *Spok2 Spok3* |
| Wa46 | *P. anserina* | naïve | - | - | absent | - |
| Wa53 | *P. anserina* | 1 | Chr 3 | 113407 | *Spok* block | *Spok2 Spok3 Spok4* |
| Wa58 | *P. anserina* | 7 | Chr 5 L | 167459 | *Spok* block | *Spok2 Spok3 Spok4* |
| Wa63 | *P. anserina* | S | - | - | absent | *Spok2* |
| Wa87 | *P. anserina* | 1 | Chr 3 | 113425 | *Spok* block | *Spok2 Spok3 Spok4* |
| Wa100 | *P. anserina* | 8 | Chr 5 L | 127869 | *Spok* block | *Spok2 Spok4* |
| Wa137 | *P. anserina* | 9 | Chr 1 | 247510 | *Spok* block | *Spok2 Spok4* |
| S | *P. anserina* | S | - | - | absent | *Spok2* |
| Psk1xS<sub>14</sub><sup>c</sup> | *P. anserina* | 1 | Chr 3 | 113407 | *Spok* block | *Spok2 Spok3 Spok4* |
| Psk7xS<sub>14</sub><sup>c</sup> | *P. anserina* | 7 | Chr 5 L | 167459 | *Spok* block | *Spok2 Spok3 Spok4* |
| T<sub>G</sub> | *P. anserina* | 5 | Chr 3 | 157028 | *Spok* block | *Spok3 Spok4* |
| Y | *P. anserina* | 5 | Chr 3 | 113435 | *Spok* block | *Spok3 Spok4* |
| Wa131 | *P. comata* | C1 | - | - | absent | *Spok1* |
| Wa139 | *P. comata* | C1 | Chr5 L | 38521 | degraded | *Spok1* |
| T<sub>D</sub> | *P. comata* | C1 | - | - | absent | *Spok1* |
| CBS 237.71 | *P. pauciseta* | P1 | Chr 4 | 74163 | *Spok* block | *Spok3 Spok4* |

<sup>a</sup>Notice that a given *Spok* block might have *Spok3*, *Spok4* or both, and this reflects the *Psk* designation.
<sup>b</sup>The location of the *Spok* block and *Enterprise* element of Wa139 are indicated in the “Genomic Location” column. *Spok1* and *Spok2* are not found within the *Spok* block.
<sup>c</sup>Lab strains created by 14 consecutive backcrosses of *Psk-1* or *Psk-7* strains to S. The size of the *Spok* block is assumed to match that of the parental strain.

### The *Spok* block moves via transposition

A number of mechanisms exist by which a sequence can move within a genome, including reciprocal translocation, ectopic recombination, and transposition (Mieczkowski, Lemoine, and Petes 2006). We can rule out reciprocal translocations in the case of the *Spok* block, as overall chromosome synteny is preserved (Vogan, Ament-Velásquez, et al. 2019). Ectopic recombination is often mediated by TEs, and transposition is, by definition, the process of TE mobility. In order to determine a candidate mode by which the *Spok* block moves, we examined the four unique *Spok* block insertion sites. This analysis revealed that no specific TE was present at the insertion site, but showed that six base pairs (RGGTAG) are always present and are repeated at the end of the *Spok* block (**Figure 1A and B**). This repeated sequence may represent a target site duplication (TSD), which is a hallmark of transposition mechanisms (Wicker et al. 2007). Additionally, the *Psk-1* and *Psk-9 Spok* block insertion sites constitute a partially palindromic sequence ATACYT||AGGTAG (**Figure 1B**), a characteristic of some TE target sites (Linheiro and Bergman 2012). Together, this finding supports an explanation whereby the *Spok* block is mobilized like a TE via transposition.

**Figure 1.**
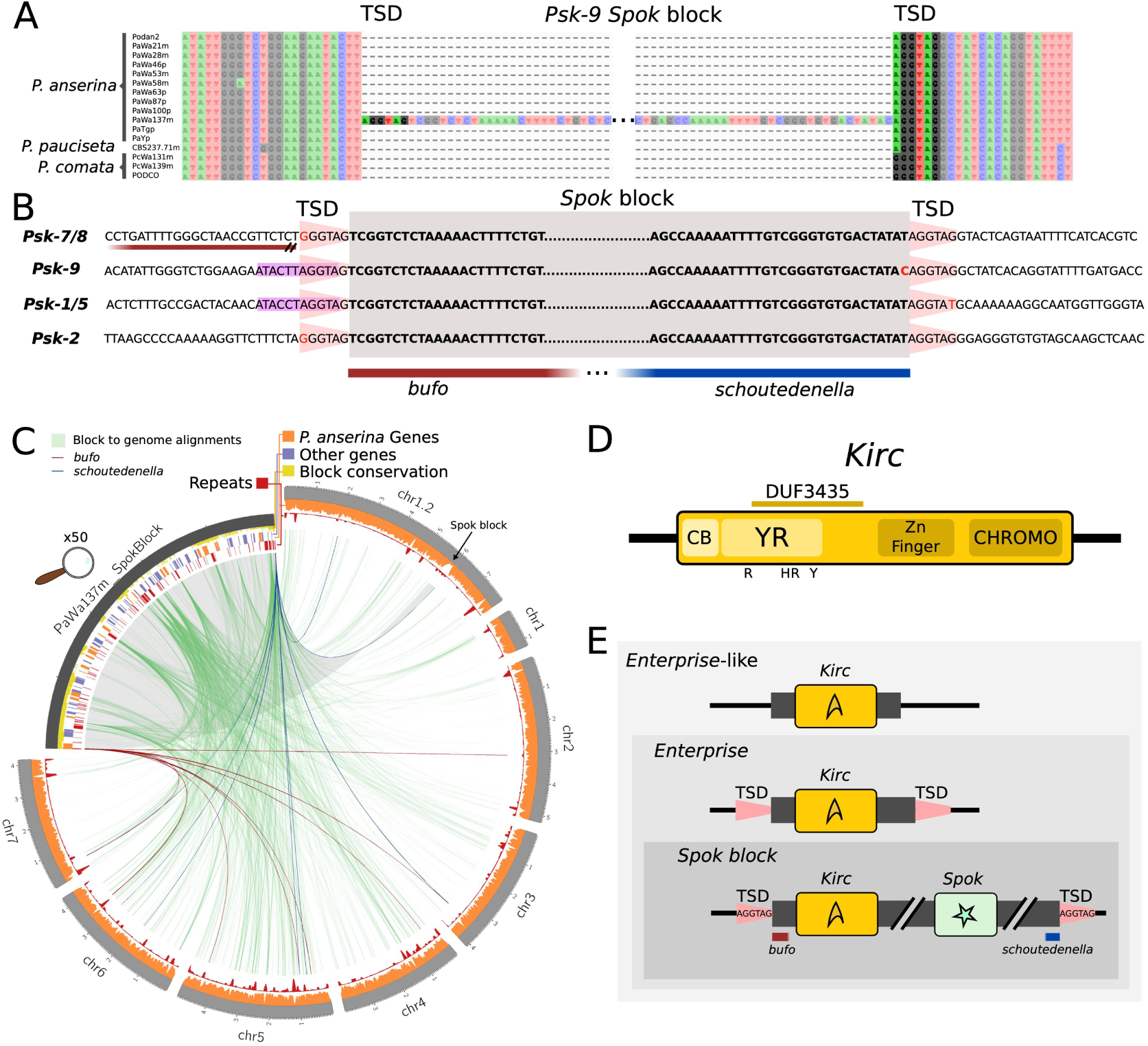
The genomic context of the *Spok* Block. **A** An alignment of the region where the *Spok* block is located in the strain Wa137 (*Psk-*9) from all *P. anserina* genomes. Putative target site duplication (TSD) is highlighted. Only the ends of the *Spok* block are shown for clarity. **B** An alignment of the ends of four versions of the *Spok* block displaying the TSD (pink trapezoid). The flanks of the *Spok* block that relate to the unclassified repeats *bufo* and *schoutedenella* are underlined in red and blue, respectively. Note that for *Psk-7*, the *Spok* block is inserted immediately next to a truncated *bufo* element. **C** A Circos plot of the genome of Wa137 (light grey track, size in Mb) aligned against its own *Spok* block (dark grey, enlarged by 50x). The tracks on the *Spok* block from outside inward represent: the conservation of regions among the different iterations of the *Spok* block where the height of the conservation track is equivalent to the number of blocks that have a given position (yellow); gene models from manual annotation of the *Spok* block, where genes with homologs within the reference genome of *P. anserina* are marked in orange and those without homologs are marked in purple; and annotated repeated elements (red). The tracks on the chromosomal scaffolds show the coverage of genic regions (orange) and repeats (red) calculated in sliding windows of 50 kb with steps of 10 kb. Green lines connect homologous segments based on MUMmer alignments. The unclassified repetitive elements *bufo* and *schoutedenella* are connected in dark red and blue segments, respectively, based on BLASTn searches. **D** Cartoon representation of the predicted protein for the ORF *Kirc* with domains predicted by bioinformatics analyes. The catalytic tetrad is marked below the protein model. **E** Cartoon model of the hierarchical nomenclature of *Enterprise*-like elements, *Enterprise* and the *Spok* block. See Supplementary Figure 3 for more detailed depictions of the *Spok* block.

The terminal sequences of TEs are often composed of structural features that are intrinsic to the transposition mechanism; LTRs for LTR retrotransposons and terminal inverted repeats (TIRs) for transposase-mobilized DNA transposons, for example (Wicker et al. 2007). These features can thus be used to determine the type of transposition or transposon underlying a given translocation. With this expectation, we examined the ends of the *Spok* block (which are very similar across all instances (Vogan, Ament-Velásquez, et al. 2019)), but found no such structural features. Curiously, the ends of the *Spok* block have been previously identified as two unclassified repetitive elements called *bufo* and *schoutedenella* (Espagne et al. 2008) (**Figure 1B**). We suspect that these unrelated elements actually represent the two ends of the *Spok* block (*bufo* representing the first 463 bp and *schoutedenella* representing the last 231 bp), and are found on their own in the genome as remains of previous, partially deleted, *Spok* block copies. To test this assumption, we mapped the location of all *bufo* and *schoutedenella* elements present in the *Podospora* genomes. We found that both elements can be identified throughout the chromosomes, with a tendency to locate in TE-rich areas in all species (**Figure 1C and Supplementary Figure 1**). In agreement with the postulate that the genomic copies of *bufo* and *schoutedenella* represent past *Spok* block insertions, at the *Psk-7* insertion site there is ∼1200 bp of sequence that is homologous to the beginning of the *Spok* block. This is nearly 700 bp longer than what *bufo* alone represents, and further implies that the *Psk-7 Spok* block was inserted at the same site as a previous, now largely deleted copy. Together, these findings further support the idea that the *Spok* block inserts at the specific target site sequence RGGTAG which is duplicated during transposition (**Figure 1B**).

### The *Spok* block accumulates sequence from the genome

As there were no structural features to guide us in our attempt to determine the mechanism through which the *Spok* block translocates, we examined the composition of the *Spok* block to identify candidate genes capable of transposing it. We annotated all genome assemblies using a modified version of the annotation pipeline in (Vogan, Ament-Velásquez, et al. 2019), relying on additional RNA-seq data, as well as an improved, manually curated repeat library (see Methods). The content of the different copies of the *Spok* block is largely overlapping (Vogan, Lorena Ament-Velásquez, et al. 2019), although the Wa137 *Spok* block has a large region of unique sequence resulting in a total size of 247 kb (**Figure 1C and Supplementary Figure 2**). Very few genes were predicted within the *Spok* block by this methodology (as evidenced by the dip in the orange track on the chr 1.2 scaffold; **Figure 1C**), so we manually annotated the *Spok* block of representative strains (Wa53, Wa28, Wa58, and Wa137), which identified numerous hypothetical protein-coding genes (e.g., 67 in the Wa137 *Spok block*), many of which appear to be pseudogenes or gene fragments. The blocks do not generally exhibit unusually high TE load. For example, the *Spok* block in Wa137 is only composed of 10.5% annotated TEs (compared to genome wide estimates of 3 – 6%, **Supplementary Table S1**), nor do they appear to have a strong signature of repeat-induced point mutation (RIP). RIP is a process that operates in numerous fungi, including *Podospora* (Graïa et al. 2001), by specifically inducing C-to-T mutations of any repeated sequence within the genome (Selker et al. 1987; Cambareri et al. 1989) and results in a drop of GC proportion in repetitive elements. Such a GC-drop is clear in many regions of the *Podospora* genome, but is conspicuously absent within the *Spok* block (Vogan, Ament-Velásquez, et al. 2019). While some of the predicted genes had identifiable homologs within the *P. anserina* reference genome (orange features in **Figure 1C**), many are absent from the reference but have sequence similarity to genes from other fungi (purple features in **Figure 1C**). Of note, many of the predicted genes have potential roles in various metabolic pathways, like metal tolerance or antimicrobial resistance (**Supplementary Table S2**). It thus seems likely that the *Spok* block has grown to its large size, at least in part, by accumulating non-repetitive sequence from elsewhere within the genome. The accumulation does not appear to have been a single event, but was more likely many individual events, as the genomic homologs are found on all chromosomes (green links in **Figure 1C**).

### The *Enterprise*

The gene annotation provided us with very few clues as to the agent(s) responsible for the translocation of the *Spok* block, so we turned to another approach. We hypothesize that the *Spok* block was a much smaller element in the past, and that such an element could be still transposing in the genomes of *Podospora* species. The closely related species *P. comata* possesses both *bufo* and *schoutedenella* repeats, and appears to have large areas of homology to the *Spok* block (**Supplementary Figure 1B**). We discovered that the *P. comata* strain Wa139 contains a 39 kb subtelomeric region on chromosome 5 (henceforth referred to as the *Enterprise*) that is nearly completely composed of TEs and sequence homologous to the *Spok* block (**Supplementary Figure 2**). The ends of the *Enterprise* are consistent with the *Spok* block (both *bufo* and *schoutedenella* are present) and it possesses the RGGTAG target site duplication (**Supplementary Figure 3A**). *Enterprise* is absent at the orthologous location in *P. anserina* and *P. pauciseta*, is much smaller in the other *P. comata* strain T_D_ (5.5 kb), and unassembled in Wa131. The other *P. comata* strains possess *bufo* and *schoutedenella* at this location, suggesting that this specific *Enterprise* insertion is now polymorphic for its state of degradation. However, it is difficult to fully recapitulate the history of the region due to the limited number of strains and the fact that Wa139 also has another *schoutedenella* copy at the insertion site that is absent in the other strains (**Supplementary Figure 3A**). Manual annotation revealed five putative genes within the Wa139 *Enterprise*, four of which are homologous to the genes within the *Spok* block and thus represent good candidates for involvement in the transposition of *Enterprise* and therefore the *Spok* block.

None of these four genes have a known function and none are *Spok* homologs. We examined their predicted protein domains, searched the *P. anserina* genomes for homologs, and scanned genomic databases for similar sequences in an attempt to discern whether they may be integral to the movement of the *Enterprise* (**Supplementary Table 3**). Only one of the genes is a likely candidate for enacting the transposition of the *Enterprise*. This gene is always present as the first ORF in the *Spok* block, but is degraded and interrupted by multiple TEs in Wa139. The ORF possesses multiple domains which may be characteristic of specific transposons, as determined by bioinformatic analysis (see Methods): a domain of unknown function called DUF3435, a zinc finger domain, and a predicted CHROMO domain (**Figure 1D**). Zinc finger domains are important for DNA binding, and CHROMO domains are implicated in histone binding and may allow TEs to be targeted to specific regions of the genome (Kordis 2005). There is strong overlap between the DUF3435 domain and tyrosine recombinases (YRs), a group of enzymes important for DNA integration of other selfish genetic elements known as *Cryptons*, parasitic plasmids, and bacteria (Kojima and Jurka 2011). YRs are extremely divergent, and there is little conserved sequence identity between this gene and known YRs, but importantly, YRs are known to have a catalytic tetrad consisting of an R-H-R-Y motif (Esposito and Scocca 1997), which is overlapping with the DUF3435 domain (**Figure 1D & Supplementary Figure 4**). Given these results, we consider this gene to be a type of YR which is responsible for transposing *Enterprise*, and name it “spore **ki**lling **r**elated **c**rypton” or *Kirc*.

From the evidence presented here, we propose that the *Spok* block and the homologous region of Wa139 represent copies of a previously unknown group of DNA transposons, which we name *Enterprise. Cryptons* are a type of DNA transposon defined only by the presence of a YR domain (Wicker et al. 2007) and may possess a TSD or not (Kojima and Jurka 2011), thus *Enterprise* can be classified as a novel group of *Crypton.* We define *Enterprise* as being composed of a YR-encoding gene homologous to *Kirc*, and possessing a TSD (**Figure 1E**). The *Spok* block therefore represents a specific version of *Enterprise* that contains *Spok* genes.

To confirm our hypothesis that *Enterprise* is capable of transposition and selfish replication, we mined fungal genomes available on JGI MycoCosm for homologous proteins of *Kirc* that are present in multiple copies within a single genome. We identified such a case in *Melanconium sp.* NRRL 54901. In this genome, a homolog of *Kirc* is found at the beginning of a ∼70 kb region that is present in four copies in the genome. Critically, this region is flanked by the same TSD as the *Spok* block and the target site is in its full palindromic context CTACCT||AGGTAG (**Supplementary Figure 5**). The only gene homologous with the *Enterprise* in *Podospora* is the *Kirc* homolog, thereby confirming that the minimal feature for transposition is the putative YR-encoding gene, *Kirc*, and classifying this region of *Melanconium sp.* as an *Enterprise.*

### *Kirc* is wide-spread in filamentous fungi

Given the fact that *Enterprise* is present in at least one species outside of *Podospora*, we queried GenBank with the sequence of *Kirc* using BLASTp to determine how widespread *Enterprise* is. We recovered a total of 481 protein hits, which were almost exclusively from within the Pezizomycotina, although putative homologs were identified from ten basidiomycete genomes as well. A phylogeny of a representative set of sequences shows that the relationships among the homologs do not follow the expected species phylogeny (**Figure 2**). Numerous species have multiple *Kirc* homologs in the same genome, some of which are not closely related. For example, within *P. anserina*, one homolog was recovered, Pa_5_10116 (see Espagne et al. (2008) for *Podospora* gene notation), which appears to be distantly related to *Kirc*, yet highly similar to homologs from *Fusarium*. Furthermore, Pa_5_10116 is pseudogenized and absent in the close relatives of *P. anserina, P. comata* and *P. pauciseta*, consistent with it being a transposable element. Attempts to describe TSDs in other fungi were largely unsuccessful as conserved flanks could not be identified in most cases, which is not unusual for *Cryptons* in general (Kojima and Jurka 2011). Thus, in the absence of additional evidence from these fungal species, we consider these as *Enterprise*-like elements. Regardless, these results suggest that *Enterprise* is a YR-mobilized group of DNA transposons that is spread throughout fungal genomes.

**Figure 2.**
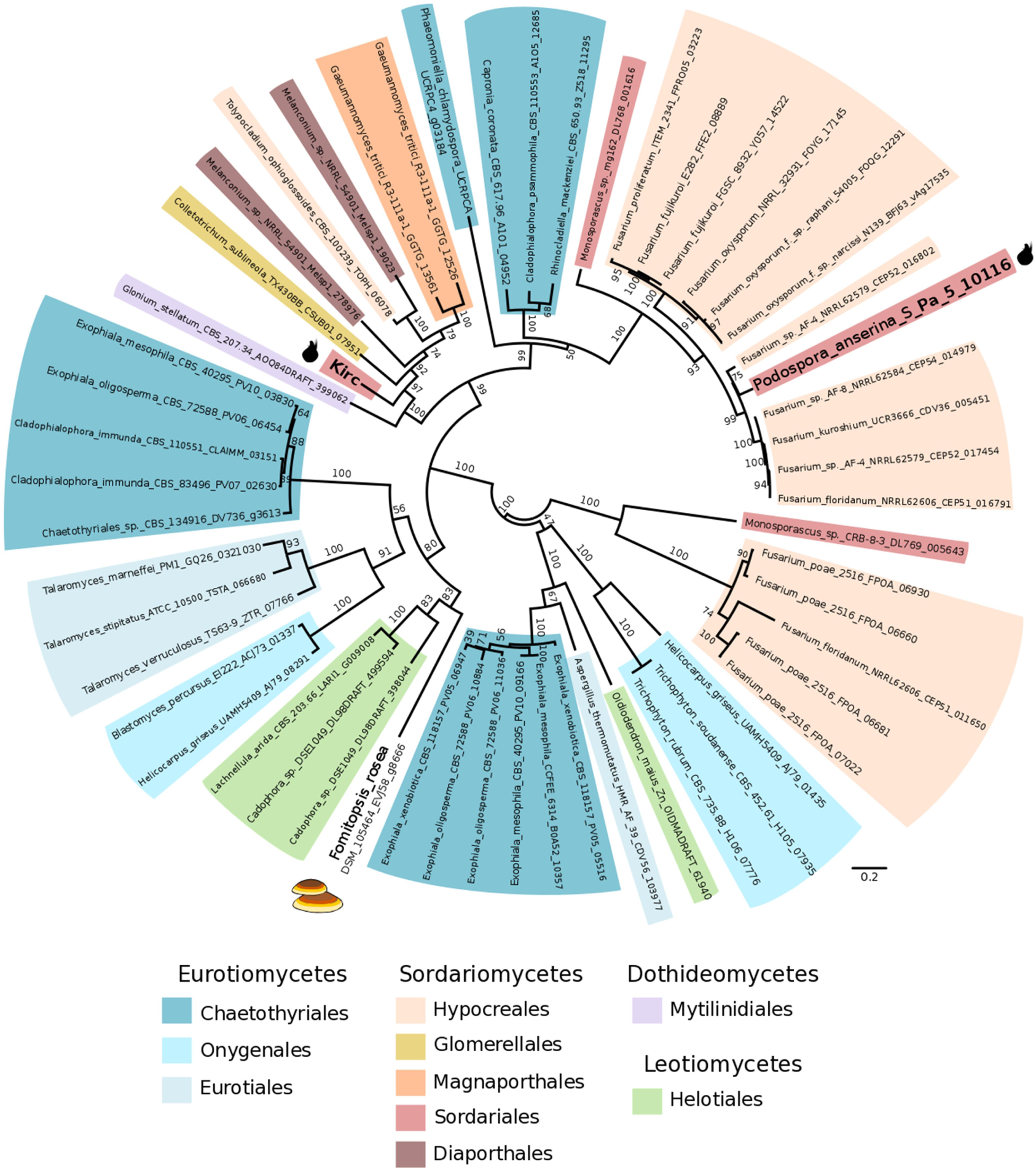
Unrooted Maximum likelihood phylogenetic tree showing the relationship between *Kirc* and other homologs from fungi. Bootstrap support values are shown above branches. Branch lengths are proportional to the scale bar (amino acid substitution per site). Taxonomic rank is indicated with coloured highlights. The sequences present in *Podospora anserina* and in the basidiomycete *Fomitopsis rosea* are marked with cartoons of the corresponding fruiting bodies. Nomenclature is formatted as species_strain_protein code.

### The *Spok* block can be deleterious

In *P. anserina*, wild strains with more than one full copy of the *Spok* block have never been found, although it is easy to generate them in the lab via crosses (Vogan, Ament-Velásquez, et al. 2019). It is possible that the burden imposed by the *Spok* block on the host is quite high, leading to selection to purge most copies in nature. To evaluate this hypothesis, we made use of backcrosses of two of the *Psk* strains (Vogan, Ament-Velásquez, et al. 2019). The backcrossed strains Psk1xS_14_ and Psk7xS_14_ are isogenic with the reference strain S, except that S has no *Spok* block. Psk1xS_14_ has the *Spok* block on chromosome 3 and induces killing in 90% of meioses, and Psk7xS_14_ has the *Spok* block on the left arm of chromosome 5 and induces spore killing in 50% of meioses. We crossed strains either to themselves (no spore killing) or to strain S (spore killing) and evaluated a number of traits related to fitness. Radial growth and time to germination of spores produced by the matings showed no variation between crosses (data not shown), however significant differences were observed among the amount of spores produced by a cross (**Figure 3**). Specifically, crosses from selfings of Psk1xS_14_ produced significantly fewer spores than from selfings of either S or Psk7xS_14_, despite the fact that no spore killing occurs, indicating that the *Psk-1 Spok* block inhibits spore production. This effect was even greater in crosses between S and Psk1xS_14_, and was most prominent when Psk1xS_14_ was used as the female, suggesting a maternal effect. With both Psk1xS_14_ and Psk7xS_14_ more spores were produced in the killing crosses than expected given the proportion of killing per ascus (Vogan, Ament-Velásquez, et al. 2019). As there were no significant differences in the amount of spores produced by Psk7xS_14_ than by S, this result suggests that the general presence of the *Spok* block itself is not deleterious, but rather that the negative effect is due to the specific content and/or to the genomic location of a given *Spok* block.

**Figure 3.**
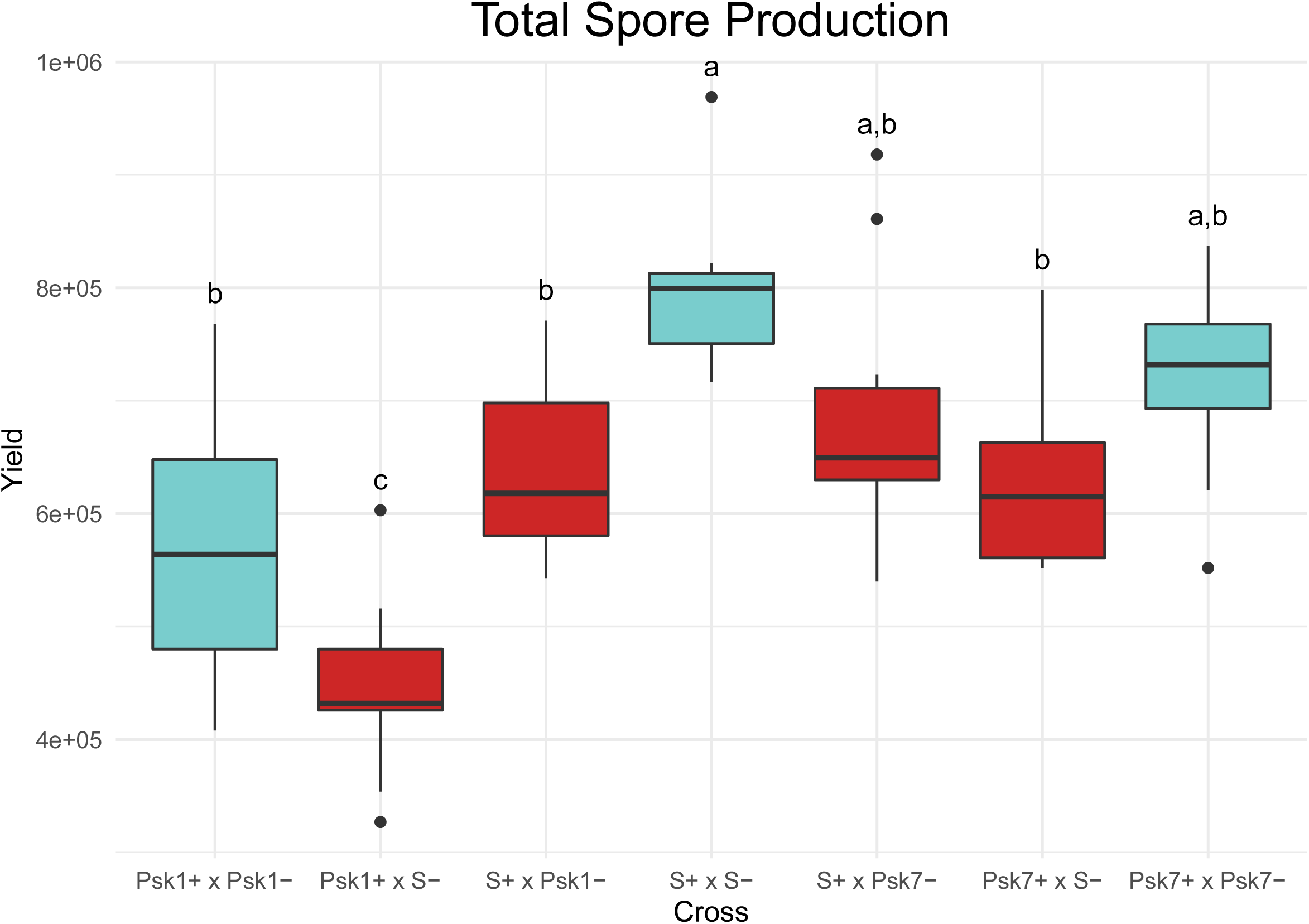
Plot showing the total amount of ascospores collected from crosses of isogenic strains possessing various iterations of the *Spok* block. Yield obtained in crosses where spore killing occurs are shown in blue; crosses with no spore killing are in red. Letters mark results of a Tukeys HSD test. The mating type is indicated by a + or – after the strain name. The *mat+* parent was always used as the maternal strain in crosses.

## Discussion

Here we provide evidence that the *Spok* block is a variant of a newly described TE, *Enterprise*, that moves throughout the genome of *P. anserina* by means of YR-mediated transposition. In addition to this ability to move, the *Spok* block is also capable of meiotic drive due to the presence of the *Spok* genes. Given the union of these two selfish properties, it can be asked who is parasitizing whom. One possibility is that *Enterprise* has hijacked the *Spoks* in order to increase its rate of transmission, thereby parasitizing a resident genomic parasite and becoming a genomic hyperparasite. Support for this commandeering can be found in the distribution of the *Spok* genes. In both *P. anserina* and *P. comata*, the *Spok* genes that are not found in association with the *Spok* block are at high frequency (*Spok2* is found in 98% of strains isolated in Wageningen (Vogan, Ament-Velásquez, et al. 2019), and so far *Spok1* has been found in all analysed strains of *P. comata*). The *Spok* block is comparatively rare (∼18% of strains from Wageningen (Vogan, Ament-Velásquez, et al. 2019)), indicating that it may prevent the *Spoks* from reaching high frequencies. Alternatively, the *Spok* genes may generally benefit from moving throughout the genome. *Spok1* and *Spok2* are found at different locations in the genome and are surrounded by TEs, suggesting they may have moved through other mechanisms like TE-mediated ectopic recombination (Vogan, Ament-Velásquez, et al. 2019). It may thus be advantageous for the *Spoks* to mobilize within TEs, like *Enterprise*, in order to change their genomic position on a regular basis, due to the fact that this relocation will result in a novel spore killing phenotype. Presumably, the population dynamics of meiotic drive ultimately decide the fate of the *Spoks* (Nauta and Hoekstra 1993, Martinossi-Allibert, et al. 2020), but the confederation of the *Spoks* and *Enterprise* as the *Spok* block may fundamentally change how effective selection is at controlling either element.

Given that the results of the fitness experiments suggest that the *Spok* block can be deleterious, there may be strong selection to purge any copies of *Enterprise*, but this may ultimately be dependent on genomic context as insinuated by the differences in phenotypic effect of carrying the *Psk-1* and *Psk-7 Spok* blocks. The *Psk-7 Spok* block is larger by nearly 50 kb, some of which includes known retrotransposons. Yet, only *Psk-1* showed a significant decrease in the amount of spores produced. It thus seems probable that the location of the block has a stronger deleterious effect rather than its content or size. The *Psk-1 Spok* block is located close to the centromere on the left arm of chromosome 3. It is possible that the increased amount of killing in *Psk-1* results in its poorer performance, although this cannot explain the observed maternal effect. The rDNA cluster resides on the same chromosome arm as the *Psk-1* block (Espagne et al. 2008). Given that this arm is only ∼700 kb and the block itself is 113 kb, the *Spok* block insertion might interfere with recombination, which is necessary for proper segregation of rDNA (Tomson et al. 2006) and likely inhibits the ability of the strain to produce viable spores. Moreover, the observation that there is not a large decrease in spore production for *Psk-1* or *Pks-7* strains when involved in spore killing suggests that strains are able to compensate for the lost spores in some way. This latter result has significant implications to our understanding of spore killing as meiotic drive, as it shifts the system away from providing the killer genotype a relative fitness advantage to an absolute fitness advantage (Nauta and Hoekstra 1993; Lyttle 1991; Martinossi-Allibert et al. 2020). Explicitly, in the naïve expectation, a spore killer that is 100% efficient would reduce the total amount of spores produced in a killing cross to half. Thus, it does not produce more total offspring with its genotype than a non-spore killing gene (relative advantage). However, if the strain carrying the spore killer is able to compensate for the loss of spore production, and produce the same or similar number of spores as in a non-killing cross (as observed here), the spore killer observes an absolute advantage, as is the case for other types of meiotic drive, like female meiotic drive. Therefore, spore killers may be more successful at invading and driving through populations than previously thought.

The *Enterprise* clearly has the ability to move large amounts of genetic material around the genome and TEs are known to be agents of horizontal gene transfer (HGT) (Schaack, Gilbert, and Feschotte 2010; Gilbert and Feschotte 2018). As such, it is plausible that *Enterprise* and related YR-mobilized DNA transposons may jump between species, and thereby transport any additional genes they may carry. The phylogeny of the *Kirc* homologs is indicative of HGT as it shows closely related *Kirc* homologs distributed amongst unrelated fungi. In at least two cases, fungal TEs have been implicated as the vehicles for gene mobility of adaptive genes. In the first case, a hAT element is associated with the HGT of toxin genes among cereal pathogens (McDonald et al. 2019). The second case comes from a recent publication which described a TE named *HEPHAESTUS* in the fungus *Paecilomyces* (Urquhart et al. 2020). It carries multiple genes that provide resistance to at least five different heavy metals and shows evidence of transfer with a distantly related species of *Penicillium*. In the latter example the TE responsible for enacting the transposition is unidentified, but strikingly, a gene with a DUF3435 domain is present at the start of *HEPHAESTUS* (annotated as *hhpA*). We find little similarity between *Kirc* and *hhpA* at the protein level, but our analyses here strongly suggest that *hhpA* is also a YR and may be responsible for the reported transposition. Whether the *Enterprise* itself can also play a role in adaptive HGT is unknown as of yet, but the potential certainly exists. We note that most of the fungal genomes in which HGT has been reported (Wisecaver and Rokas 2015) possess genes containing DUF3435 domains, which together with the data from *HEPHAESTUS* suggests a convergent role in this type of TE as a vector for HGT of adaptive genes in fungi.

## Conclusions

The constant ‘tug-of-war’ between TEs linking themselves to host genes, and the actions of genome defense and selection to purge them is of key importance to the evolution of genome architecture. It is likely that we are witnessing this fight play out to the extreme in *P. anserina*, with the high effectiveness of both RIP and the *Spok* block, making *Podospora* an ideal system in which to continue to study genomic conflict. The discovery that the up to 247 kb large *Spok* block likely transposes through a YR-mediated mechanism moves the upper limit of TE size by nearly 100 kb, and its hyper-selfishness combining TE mobility and meiotic drive adds a new dynamic by which selfish elements exploit their host genome. This study not only changes our perception of how TEs influence genome evolution, but also broadens the horizons in terms of what may be possible through genetic manipulations in the laboratory. Understanding the molecular mechanism of YR-mediated transposition should thus be of prime interest in future research.

## Methods

### Fungal material

Strains used in this study were obtained from the Wageningen collection and cultivated as in Vogan, Ament-Velásquez, et al. (2019). Strain S was used as the standard reference strain with no *Spok* block. As spore killer strains for the fitness experiments (see below) we used the backcrossed strains Psk1xS_14_ and Psk7xS_14_ (Vogan, Ament-Velásquez, et al. 2019), which should be isogenic to S but with a *Spok* block in chromosome 3 and 5, respectively. The more recently isolated strains Wa131, Wa137 and Wa139 were sampled during the fall of 2016 around Wageningen (the Netherlands) from dung of rabbit (Wa131 and Wa137, locality Unksepad Oosterbeek) or horse (Wa139, locality Uiterwaarden Wolfswaard). Morphological differences like smaller perithecia and abundant tomentose apricot-colored mycelium in HPM medium (Vogan, Ament-Velásquez, et al. 2019), as well as analyses of sequence data, allowed us to assign Wa131 and Wa139 to the species *P. comata*. Previously, only one strain from this species, T_D_, was known (Boucher, Nguyen, and Silar 2017; Vogan, Ament-Velásquez, et al. 2019), hence these new strains constitute a new report of this species for the Netherlands.

### DNA and RNA extraction and sequencing

Following Vogan et al. (Vogan, Ament-Velásquez, et al. 2019), we grew monokaryotic strains on PASM0.2 plates covered with a layer of cellophane. Genomic DNA for short-read sequencing was extracted from 80-100 mg of fungal tissue with the Fungal/Bacterial Microprep kit (Zymo; www.zymo.com). Paired-end libraries were prepared and sequenced using the Illumina HiSeq X (150-bp-long) technology at the SNP and SEQ Technology platform (SciLifeLab, Uppsala, Sweden). For RNA extraction, around 150 mg of harvested mycelium were frozen in liquid nitrogen and stored at -80°C. We extracted total RNA from the grounded frozen tissue using the RNeasy Plant Mini Kit (Qiagen, Hilden, Germany). Quality was checked on the agilent 2100 Bioanalyzer (Agilent Technologies, USA) and the RNA was treated with DNasel (Thermo Scientific). The sequencing library was prepared with a NEBNext Ultra Directional RNA Library Prep Kit for Illumina (New England Biolabs). We purified polyA+ transcripts with the NEBNext Poly(A) mRNA Magnetic Isolation Module (New England Biolabs). A paired-end library was sequenced with Illumina HiSeq 2500 at the SNP and SEQ Technology platform.

For long-read sequencing, we grew the monokaryotic strains in PASM0.2 plates, from where we sliced small agar cubes to inoculate liquid cultures of 200 ml 3% malt extract solution, which were subsequently incubated in a shaker for 10-14 days at 27°C (Vogan, Ament-Velásquez, et al. 2019). Mycelium aggregates were filtered from the flasks, any remaining agar was removed, and around 1 g was stored at -20°C. As described in Sun et al. (2017), the tissue was freeze-dried and macerated, followed by DNA extraction using Genomic Tip G-500 columns (Qiagen) and cleaning with the PowerClean DNA Clean-Up kit (MoBio Labs). Additionally, DNA was purified using magnetic beads (Speed-Beads, GE) and eluted for 20 min at 37°C followed by overnight storage at 4°C twice to increase concentration (around 65 ng/μl). Wa137-was sequenced on an R9.5.1 Flowcell (Oxford Nanopore Technologies) with a modified SQK-RAD004 protocol using 550 ng DNA to 1.5μl FRA to increase read lengths. Wa139-was prepared using the ligation protocol (SQK-LSK109) and sequenced on an R9.4.1 flowcell. Basecalling was done using Guppy v. 1.6.

### Genome assembly

For most strains we used the assemblies produced in Vogan & Ament-Velásquez et al. (2019). For newly sequenced strains, we produced new assemblies as follows. The adapters from the Illumina HiSeq reads were identified with cutadapt v. 1.13 (Martin 2011) and removed using Trimmomatic v. 0.36 (Bolger, Lohse, and Usadel 2014) using the following options: ILLUMINACLIP:adapters.fasta:1:30:9 LEADING:20 TRAILING:20 SLIDINGWINDOW:4:20 MINLEN:30. Pairs with both forward and reverse reads after filtering were used for downstream analyses. For the strain Wa131, which only has Illumina data, we used SPAdes v. 3.12.0 (Bankevich et al. 2012) with the k-mers 21,33,55,77 and the *--careful* option. For the strains Wa137 and Wa139, the MinION reads with a mean Phred quality (QV) above 9 and longer than 1 kb were assembled using Minimap2 v. 2.11 and Miniasm v. 0.2 (Li 2018, 2016). The resulting assembly was polished twice with Racon v. 1.3.1 (Vaser et al. 2017) using all MinION reads (no filtering). Further polishing was done with the filtered Illumina reads in five consecutive rounds of Pilon v. 1.22 (Walker et al. 2014). We used BWA v. 0.7.17 (Li and Durbin 2010) for short-read mapping, with PCR duplicates marked using Picard v. 2.18.11 (http://broadinstitute.github.io/picard/), as well as local indel re-alignment using the Genome Analysis Toolkit (GATK) v. 3.7 (Van der Auwera et al. 2013).

We assigned the scaffolds to chromosomes based on alignments to the reference genome of the S strain (Espagne et al. 2008), available at the Joint Genome Institute MycoCosm website (https://mycocosm.jgi.doe.gov/mycocosm/home) as “Podan2” (Grigoriev et al. 2014). We discarded small contigs (<100kb) of rDNA repeats as well as mitochondrial-derived sequences, except for the largest mitochondrial contig. We assessed the quality of the final assemblies by visual inspection of the mapping of both long and short reads using Minimap2 and BWA, respectively. Mean depth of coverage was calculated with QualiMap v.2.2 (Okonechnikov, Conesa, and García-Alcalde 2016). Other assembly statistics were calculated with QUAST v. 4.6.3 (Mikheenko et al. 2016).

### Genome annotation

A GitHub repository is available with Snakemake v. 5.4.4 (Köster and Rahmann 2012) pipelines relevant to genome annotation at https://github.com/johannessonlab/SpokBlockPaper.

The TEs and other repeats in *P. anserina* were classified previously by Espagne et al. (2008) based on the original reference genome of the S strain or “Podan1”, and is hereafter referred to as the “Espagne library”. To explore the diversity of TEs in the newly generated *Podospora* genomes, we identified repeats de novo and manually compared them to the Espagne library to identify duplicates and new elements. Specifically, we ran RepeatModeler v. 1.0.8 (http://www.repeatmasker.org/RepeatModeler/) on the scaffolds larger than 50 kb of all available long-read assemblies (Snakemake pipeline *PaTEs.smk*). Each resulting RepeatModeler consensus was BLASTn-searched back to the original genome and the best 20 hits with 2-kb flanks were aligned with T-Coffee v. 12.00.7fb08c2 (Notredame, Higgins, and Heringa 2000) (*TEManualCuration.smk*), and visually inspected for manual curation. The curated consensuses were assigned to the Espagne et al. (2008) equivalents based on similarity (allowing for RIP-induced mutations) or were given a new name when having no homology to anything in the Espagne library. It was discovered that the *gypsy* element *crapaud* has numerous diverged copies with unique LTRs. We annotated all *crapaud* LTRs that were in multiple copies within *P. anserina* individually to improve repeat masking. We refer to the final repeat library as “PodoTE-1.00” (available at the GitHub repository).

To generate a genome annotation of all assemblies, we ran an updated version of the pipeline in Vogan & Ament-Velásquez et al. (2019), named *PaAnnotation.smk*. We used MAKER v. 3.01.2 (Holt and Yandell 2011; Campbell et al. 2014) with the previously produced training files used for the ab initio prediction programs GeneMark-ES v. 4.32 (Lomsadze et al. 2005; Ter-Hovhannisyan et al. 2008) and SNAP release 2013-06-16 (Lomsadze et al. 2005), as well as the following dependencies: RepeatMasker v. 4.0.7 (http://www.repeatmasker.org/), BLAST suite 2.6.0+ (Camacho et al. 2009), Exonerate v. 2.2.0 (Slater and Birney 2005), and tRNAscan-SE v. 1.3.1 (Lowe and Eddy 1997). As evidence, we used STAR v. 2.6.1b (Dobin et al. 2013) to produce transcript models (maximum intron length set to 1000 bp) of various RNA-seq data sets. Specifically, we mapped the reads of the monokaryotic isolate Wa63- (*P. anserina*) to the assembly PaWa63m (Vogan, Ament-Velásquez, et al. 2019), of the monokaryotic isolate Wa131- (*P. comata*) to the assembly PcWa139m (this study), and of the dikaryotic Psk7xS_14_ (*P. anserina*) to the assembly PaWa58m (Vogan, Ament-Velásquez, et al. 2019). We then processed the mapped reads with Cufflinks v. 2.2.1 (Trapnell et al. 2010) to obtain the transcript models. As external evidence, we used CDS from the Podan2 annotation, protein sequences from the T strain of *P. comata* (Silar et al. 2019), and a small dataset of manually curated proteins. To aid in manual curation of selected regions (mostly the *Spok* block), we visually inspected the mapping of RNA-seq reads of the different datasets, along with CDS produced with TransDecoder v. 5.5.0 (Haas et al. 2013) on the Cufflinks models, as well as the output of RepeatMasker ran externally from MAKER with the PodoTE-1.00 library. Additionally, we queried predicted gene models into the NCBI databases (NCBI Resource Coordinators 2016) to verify the annotations.

The *Kirc* protein sequence was analyzed with HHPred (Zimmermann et al. 2018) and Gremlin (Balakrishnan et al. 2011). The Gremlin-generated alignment of *Kirc* homologs was used to generate region-specific sequence logos with Weblogo (Crooks et al. 2004). The relationship of *Kirc* to other YRs was confirmed by comparing the sequence to the crystal structure of known YRs (CRE (PDB code 3mgv), XERD (1a0p), and FLP (1flo)) as well as the protein sequence from the transposable element *Crypton-Cn1* using the software Promals3D (Pei, Kim, and Grishin 2008).

### Genome alignments

We used the NUCmer program from the MUMmer package v. 4.0.0beta2 (Kurtz et al. 2004) using the parameters *-b 200 -c 22 --maxmatch* to align the *Spok* blocks to each other, and *changed to -c 40* for whole-genome assemblies. To achieve higher sensitivity, we used BLASTn from the BLAST suite 2.9.0 (Camacho et al. 2009) to search for the presence of the unclassified repeats *bufo* and *schoutenella*. Both the NUCmer and the BLAST alignments were plotted using Circos v. 0.69.6 (Krzywinski et al. 2009) along with manual curations of coding regions and repetitive elements. The distribution of TE and gene content along chromosomes was calculated in windows of 50 kb with steps of 10 kb using BEDtools v. 2.29.0 (Quinlan and Hall 2010; Quinlan 2014), with the utilities *makewindows* and *coverage*. The fraction of conservation between blocks compared to the block in Wa137 was calculated by aligning the block sequences (within the TSD) of Wa28 (*Psk-2*), Wa53 (*Psk-1*), Wa58 (*Psk-7*) and Wa139 with NUCmer and the BEDtools utility *genomecov*. The Snakemake pipelines used to produce the Circos plots (*CircosBlock.smk* and *CircosAllBlocks.smk*) are available at h https://github.com/johannessonlab/SpokBlockPaper.

### Phylogenetic analyses

Homologs of *Kirc* were identified from GenBank using BLASTp with the truncated version of *Kirc* from Wa53 that has no CHROMO domain. Nucleotide sequences of hits with e-values < 1e-100 were compiled along with a homolog from *P. anserina* (Pa_5_10116), and two homologs from *Melanconium sp.* NRRL 54901 extracted from MycoCosm (see below), and aligned with MACSE v. 2.03 (Ranwez et al. 2018). We used TrimAl v. 1.4.1 (Capella-Gutierrez, Silla-Martinez, and Gabaldon 2009) to trim the resulting protein alignment with the *-gappyout* option. We then used IQTree v. 1.6.8 (Kalyaanamoorthy et al. 2017; Nguyen et al. 2014) to produce a Maximum Likelihood phylogeny with extended model selection (-m MFP). To estimate the branch support, we produced 1000 standard bootstrap pseudoreplicates.

To search Mycocosm for other copies of *Enterprise* the following approach was taken. The protein sequence of *Kirc* was used as a query with BLASTX against all genomes within MycoCosm (as of February 2019). Genomes with multiple high confidence positive hits were identified and the regions with putative *Kirc* homologs were manually extracted. Priority was given to genomes where the hits were associated with large duplicated regions (>50 kb). *Melanconium sp.* NRRL 54901 (produced as part of the 1KFG project; Spatafora et al. 2011) had the most copies with clear termini.

### Fitness assays

The cultures used for the crosses were revived from the -80 freezer on PASM0.2 (van Diepeningen et al. 2008) at 27°C for several days and then stored at +4°C until use. Strains were grown for 5 days on fresh PASM0.2 plates before inoculating the cross. In the crosses, one strain was grown as mycelia and thereby assigned the female role, while a compatible strain of the other mating type was assigned the male role by fertilizing the mycelia with microconidia. The strain that was assigned the female role was grown in a 35 mm petri dish with 5 ml HPM medium (Vogan, Ament-Velásquez, et al. 2019) by inoculating a small cube of agar with mycelium (∼2 x 2 mm). In parallel, the strain that was assigned the male role was grown on a 90 mm petri dish with micro conidiation medium (Esser 1974; King 2013) by inoculating seven plugs of mycelium spread over the plate. After 7 days of growth, microconidia were harvested by adding 5 ml of sterile water to the plate and sweep over the mycelium with a drigalski spatula for 1 minute. The female mycelium was then fertilized with 0.5 ml of the microconidial suspension. The suspension was carefully spread out to make sure all mycelium was covered. The fertilized mycelia were then further incubated under standard conditions (27°C, 12/12 light/dark cycle) (Vogan, Ament-Velásquez, et al. 2019). The cultures were monitored daily for signs of spores shot from the asci in order to score the first day of spore-shooting. To reduce the complexity of the experiment, the strains used as female were always of mating type *mat+*.

At 6 days post-fertilization, single spores were collected with a needle to measure germination frequency and growth speed. From each cross, 10 spores from 4-spored asci were picked, and in cases with spore killing, an additional 10 spores from 2-spored asci were picked. The 10 spores were transferred to a single 90 mm petri dish with PASM2 medium (van Diepeningen et al. 2008) with 0.4% ammonium acetate added (to activate the spores) (Esser 1974; King 2013). Spores were spaced out in a predetermined pattern (4 lines of 2, 3, 3, 2 spores). After two days of incubation, the germination was scored and colony diameter was measured in two directions. If there was no growth microscopic inspection was performed to check whether a spore was present in the agar to avoid scoring no germination in case the inoculation failed.

At 12 days post-fertilization, spores were harvested from the lids of each crossed culture and used for estimating total spore yield. Spores were collected by pipetting 750 µl of harvest liquid (1M NAOH, 0.025% SDS) in the lid. Spores were then scraped off the lid using the pipette tip. The liquid was then collected into a 2 ml Eppendorf tube. Another 750 µl of harvest liquid was used to repeat the process to make sure most of the spores were collected from the lid. The tubes were then heated for 4 hours at 85°C, then shaken in a Qiagen Tissuelyser for 90 seconds at 30 Hz. After this, the tubes were stored at 4°C overnight. The cooled tubes were again shaken in a Qiagen Tissuelyser for 90 seconds at 30 Hz. This process prevents the clumping of spores. Total yield was determined by counting the amount of spores in a volume of 5 µl of 50x diluted suspension pipetted on an object glass using a stereomicroscope. Counts were taken five times for each replicate cross. Statistical analyses were conducted in base R v. 3.5.0 to determine significance and power.

## Acknowledgements

This work was supported by the European Research Council (ERC) grant ERC-2014-CoG (project 648143, SpoKiGen) and The Swedish Research Council to H.J, and support from the Lars Hierta Memorial Foundation and The Nilsson-Ehle Endowments of the Royal Physiographic Society of Lund to S.L.A.V. We thank the support given by the National Genomics Infrastructure (NGI) / Uppsala Genome center on massive parallel DNA sequencing. The computations were performed on resources provided by SNIC through Uppsala Multidisciplinary Center for Advanced Computational Science (UPPMAX) under the projects SNIC 2017/1-567 and SNIC 2019/8-371. The sequence data of Melanconium sp. NRRL 54901 was produced by the US Department of Energy Joint Genome Institute http://www.jgi.doe.gov/ in collaboration with the user community. We would also like to thank Sergio Tusso and the TE Jamboree of the Suh’s Lab for useful advice.

## Supplementary Figures

**Supplementary Figure 1.**
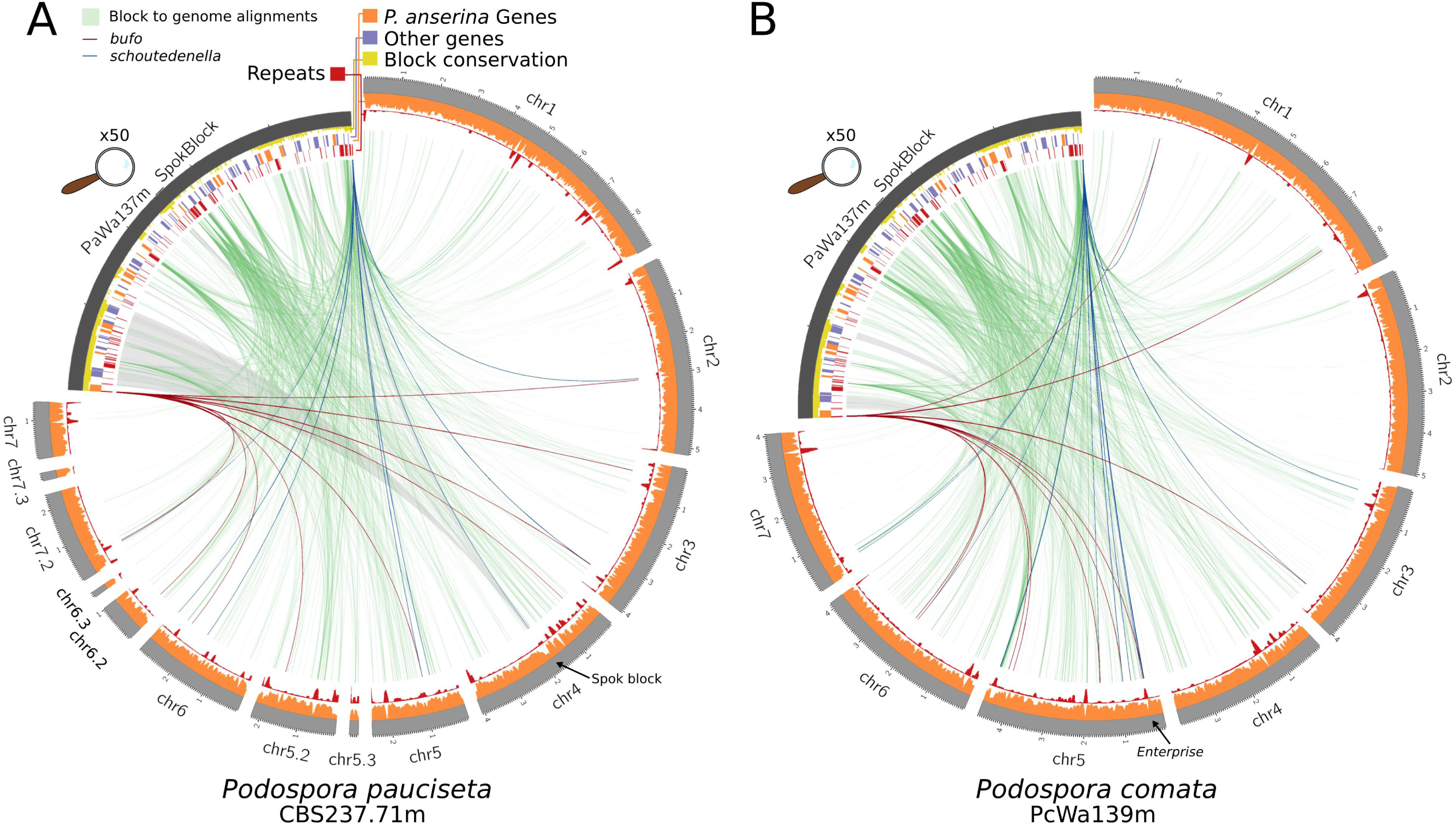
Circos plots comparing the Wa137 *Spok* block to the genome of **A** the *P. pauciseta* strain CBS237.71 and **B** the *P. comata* strain Wa139.

**Supplementary Figure 2.**
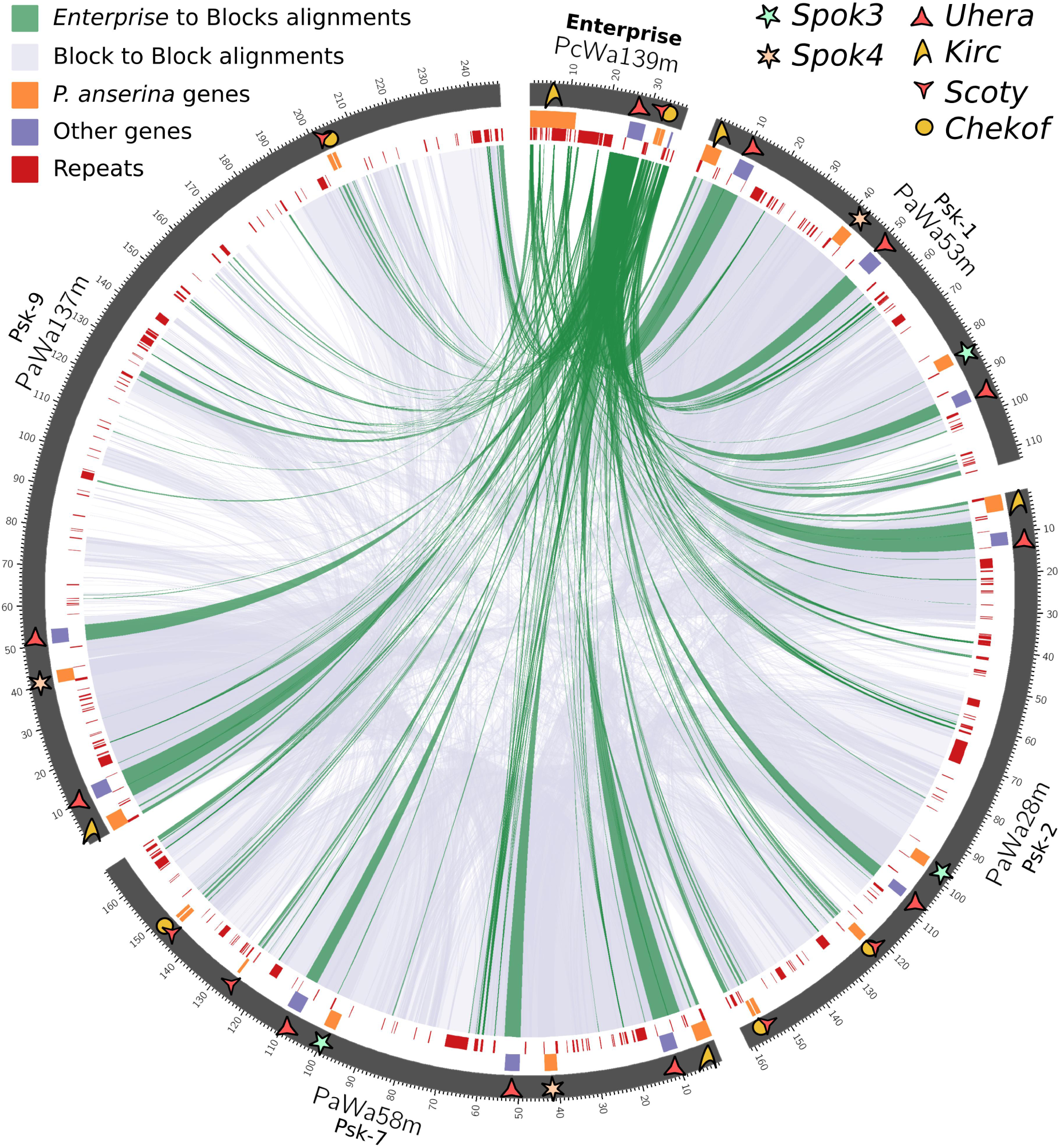
A circos plot comparing the *Enterprise* of five different strains. Four of these represent different *Spok* blocks. Dark green lines connect homologous segments of the Wa139 *Enterprise* to the various *Spok* blocks. Lilac lines show homologous regions among the *Spok* blocks. Genes of interest are marked with symbols (See Supplementary Table 3).

**Supplementary Figure 3.**
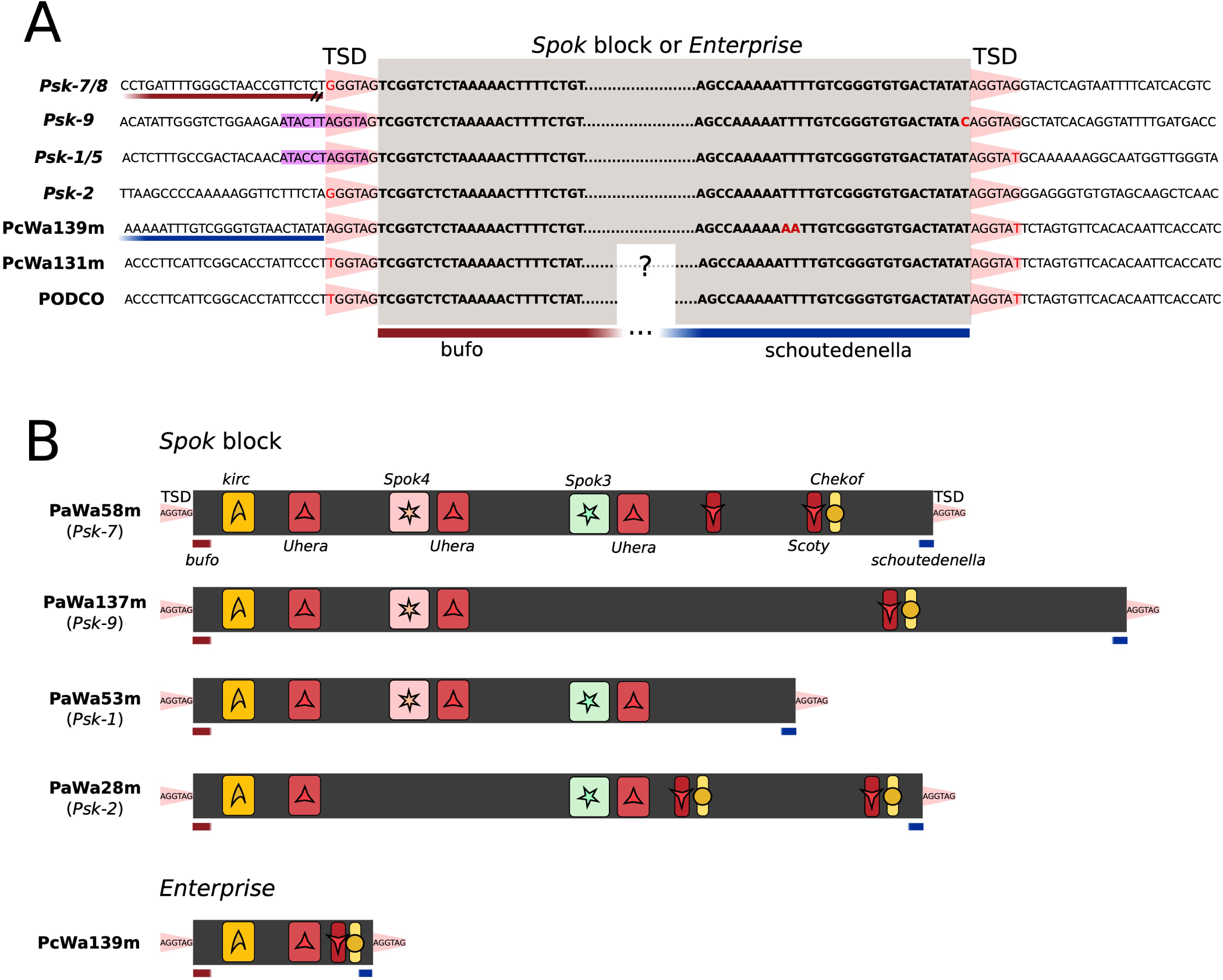
Description of the *Spok* Block. **A** An alignment of the ends of four versions of the *Spok* block displaying the TSD (red trapezoid) plus the insertion site of *Enterprise* in three *P. comata* strains. The majority of *Enterprise* is deleted in the strain T_D_ (PODCO) and unassembled in Wa131. Additionally, Wa139 is inserted next to a copy of *schoutedenella* that is absent in both PODCO and Wa131. **B** Cartoon models representing the structure and gene content of four *Spok* blocks and of *Enterprise* from Wa139. Relevant genes and features are annotated. Not to scale.

**Supplementary Figure 4.**
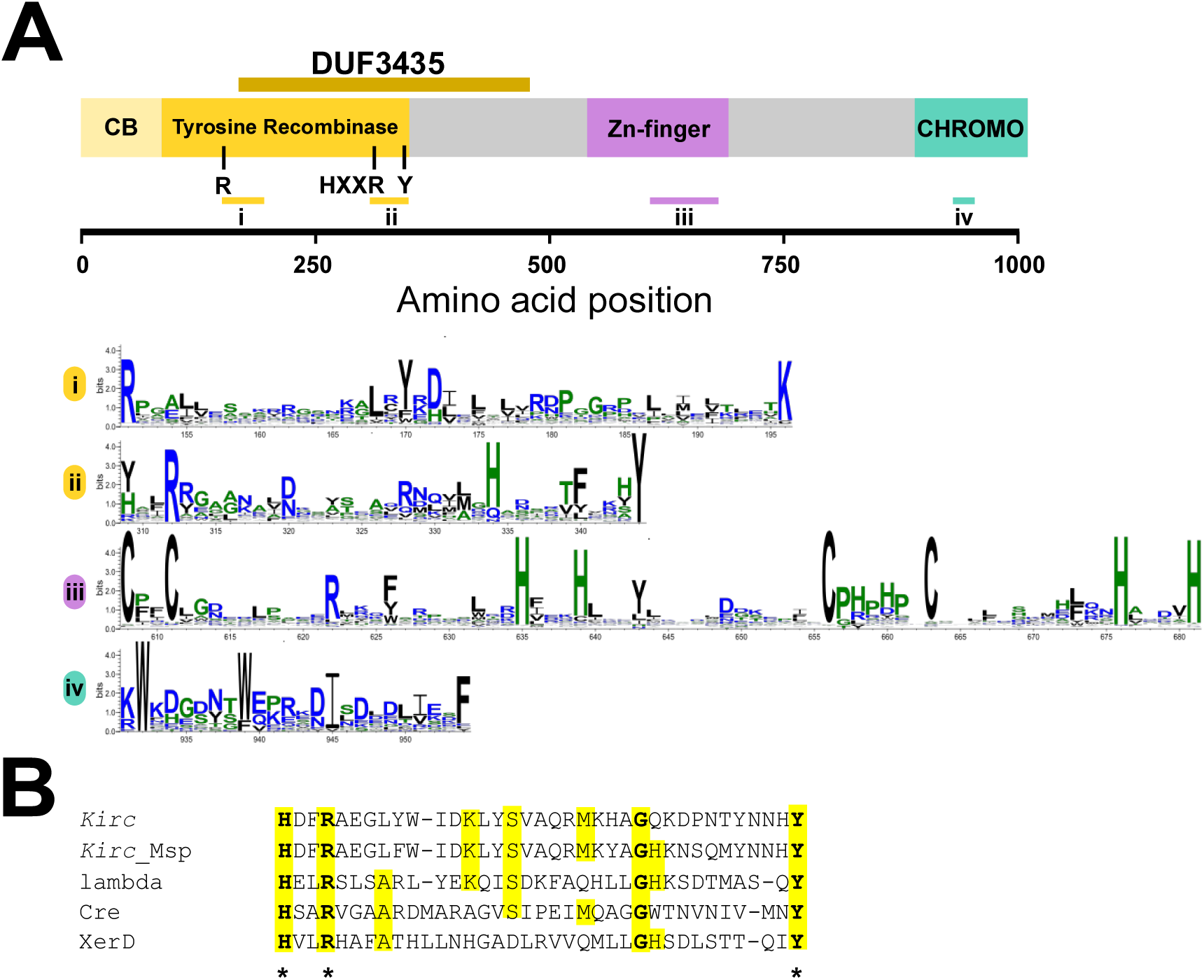
Domain annotation of the *Kirc* protein obtained using HHPred. Four domains were identified: a N-terminal tyrosine recombinase domain with the associated core-binding domain, a central Zn-finger domain and a C-terminal chromodomain. An alignment of 279 *Kirc* homologs generated by Gremlin with HHBlits was used to create sequence logo of relevant regions of the *Kirc* sequences, including the active site regions of the tyrosine recombinase domain (i and ii), the Zn-finger region (iii) and the chromodomain region (iv) using Weblogo. **B**. Alignment of the *Kirc* sequence with an active site region of known tyrosine recombinases together with the sequence of a *Kirc* homolog from *Melanconium sp.*. Alignment is based on the HHPred output. Active site residues are marked with an asterisk. Kirc: *Kirc* gene in the *Spok* block of strain Wa58 (sites 309-344); Kirc_Msp: CE55503_10527 (110-145); lambda: lambda integrase PDB: 5J0N (308-342); Cre: recombinase enterobacteriaphage P1, PDB: 1XO0 (270-305); XerD: XerD site-specific recombinase E. coli, PDB: 1A0P (242-277).

**Supplementary Figure 5.**
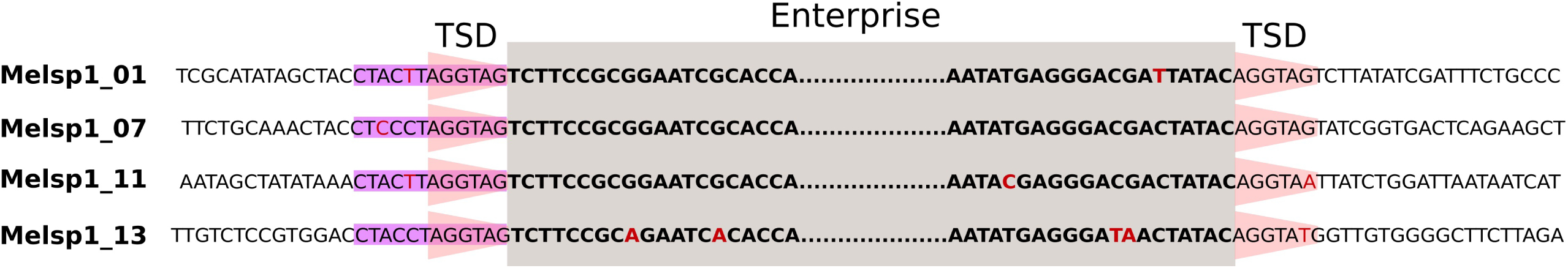
Alignment of four *Enterprise* elements within *Melanconnium sp.* revealing the TSD.

**Supplementary Table 1.** Assembly statistics for all strains with genomic data used in this study.

| Species | Strain | Assembly | Technology | Assembler | Scaffolds<sup>a</sup> | N50 | GC content | Size (bp) | Mean Depth long (x) |
| --- | --- | --- | --- | --- | --- | --- | --- | --- | --- |
| *P. anserina* | Wa21- | PaWa21m | PacBio | HGAP 3.0 | 9 | 5072363 | 51.49 | 35767411 | 80.3071 |
| *P. anserina* | Wa28- | PaWa28m | PacBio | HGAP 3.0 | 7 | 5076631 | 51.41 | 35929333 | 76.2045 |
| *P. anserina* | Wa46+ | PaWa46p | PacBio | HGAP 3.0 | 9 | 4338573 | 51.37 | 35724019 | 117.8295 |
| *P. anserina* | Wa53- | PaWa53m | PacBio | HGAP 3.0 | 7 | 4924837 | 51.52 | 35808992 | 83.821 |
| *P. anserina* | Wa58- | PaWa58m | PacBio | HGAP 3.0 | 7 | 5037662 | 51.46 | 35985748 | 112.1709 |
| *P. anserina* | Wa63+ | PaWa63p | PacBio | HGAP 3.0 | 7 | 4907271 | 51.36 | 36002190 | 111.2245 |
| *P. anserina* | Wa87+ | PaWa87p | PacBio | HGAP 3.0 | 9 | 4338609 | 51.44 | 35896992 | 105.8792 |
| *P. anserina* | Wa100+ | PaWa100p | PacBio | HGAP 3.0 | 7 | 5096663 | 51.5 | 35953417 | 114.091 |
| *P. anserina* | Wa137- | PaWa137m | MinION | Miniasm 0.2 | 9 | 4341106 | 51.08 | 36531367 | 49.924 |
| *P. anserina* | Y+ | PaYp | MinION | Miniasm 0.2 | 8 | 4821330 | 51.55 | 35719817 | 35.6112 |
| *P. anserina* | T<sub>G</sub>+ | PaTgp | MinION | Miniasm 0.2 | 13 | 4230402 | 51.54 | 35778794 | 37.6743 |
| *P. pauciseta* | CBS237.71- | CBS237.71m | MinION | Miniasm 0.2 | 13 | 4070182 | 51.6 | 35576233 | 24.3067 |
| *P. comata* | Wa131- | PcWa131m | Illumina HiSeq | SPAdes 3.12.0 | 1119 | 258199 | 52.1 | 34680523 | - |
| *P. comata* | Wa139- | PcWa139m | MinION | Miniasm 0.2 | 7 | 4603841 | 51.98 | 34819024 | 52.5779 |
| ***P. anserina*** | **S+** | **Podan2** | **WGS** | **Arachne** | **7** | **4734292** | **52.17** | **35010595** | **-** |
| ***P. comata*** | **T<sub>D</sub>+** | **PODCO** | **454 + Illumina** | **Newbler 2.3** | **7** | **4662433** | **52.42** | **34385490** | **-** |

<sup>a</sup>The number of scaffolds correspond to those that map to the 7 chromosomes (excluding mitochondrial and rDNA contigs), but other columns include all scaffolds
<sup>b</sup>Numbers correspond to long-read data when available, otherwise to short-read data. For Wa139- and Wa137- the reads were filtered for QV>9 and to be larger than 1kb, resulting in 184301 and 202279 reads, respectively.
<sup>c</sup>Scaffolds larger than 50kb, excluding the mitochondrial and rDNA scaffolds

| Species | Strain | Assembly | Mean Depth<br>short (x) | No. Reads<sup>b</sup> | Mean Read<br>Length (bp) | Repeat<br>Content (%)<sup>c</sup> | Source |
| --- | --- | --- | --- | --- | --- | --- | --- |
| *P. anserina* | Wa21- | PaWa21m | 78.0394 | 313285 | 11863 | 5.893 | Vogan et al. 2019 |
| *P. anserina* | Wa28- | PaWa28m | 81.0651 | 327199 | 10105 | 5.961 | Vogan et al. 2019 |
| *P. anserina* | Wa46+ | PaWa46p | 107.4690 | 363721 | 12949 | 6.042 | Vogan et al. 2019 |
| *P. anserina* | Wa53- | PaWa53m | 78.7383 | 327523 | 11382 | 5.924 | Vogan et al. 2019 |
| *P. anserina* | Wa58- | PaWa58m | 88.3996 | 354055 | 13130 | 6.109 | Vogan et al. 2019 |
| *P. anserina* | Wa63+ | PaWa63p | 77.4007 | 351077 | 12562 | 6.223 | Vogan et al. 2019 |
| *P. anserina* | Wa87+ | PaWa87p | 74.3914 | 356924 | 12928 | 6.171 | Vogan et al. 2019 |
| *P. anserina* | Wa100+ | PaWa100p | 80.9098 | 361645 | 12857 | 5.914 | Vogan et al. 2019 |
| *P. anserina* | Wa137- | PaWa137m | 143.6921 | 330780 | 7913 | 6.047 | This study |
| *P. anserina* | Y+ | PaYp | 83.7234 | 697535 | 1952 | 5.873 | Vogan et al. 2019 |
| *P. anserina* | T<sub>G</sub>+ | PaTgp | 155.9337 | 1286164 | 1384 | 6.009 | Vogan et al. 2019 |
| *P. pauciseta* | CBS237.71- | CBS237.71m | 105.7037 | 240014 | 6188 | 6.056 | Vogan et al. 2019 |
| *P. comata* | Wa131- | PcWa131m | 162.0216 | 39127244 | 150 | 3.555 | This study |
| *P. comata* | Wa139- | PcWa139m | 150.9355 | 236001 | 5786 | 3.769 | This study |
| ***P. anserina*** | **S+** | **Podan2** | 10 | - | - | 4.629 | Espagne et al. 2008 |
| ***P. comata*** | **T<sub>D</sub>+** | **PODCO** | 22 | - | - | 3.054 | Silar et al. 2018 |

**Supplementary Table 2.** Hypothetical genes from within the *Spok* block of Wa137. Pfam domains inferred from NCBI conserved domain database. Putative functions inferred from BLASTp to other genes with functional annotations. Blank cells indicate either no pfam hits, or genes with BLASTp results to only hypothetical genes.

| Gene code \| name in Wa137 | pfam |  |  |  | Putative function |
| --- | --- | --- | --- | --- | --- |
| gene01805 \| Kirc | DUF3435<br>superfamily | CD_CSD<br>superfamily | CHROMO |  |  |
| gene01806 \| Uhera1 | DEAD-like_<br>helicase_C<br>superfamily | DEAD-like_helicase_N<br>superfamily |  |  |  |
| PaWa137m.02400 \| PpBr36_10652_<br>Pyricularia_pennisetigena | Peptidase_C48<br>superfamily |  |  |  |  |
| PaWa137m.02401 \|<br>Pa_2_5490_homolog | NAP |  |  |  |  |
| PaWa137m.02402 \| S7711_11618_<br>Stachybotrys_chartarum_homolog |  |  |  |  | NAD-dependent protein<br>deacetylase |
| PaWa137m_chromosome_1.2:hsp:12095<br>1:4.5.0.58 \| Pa_7_10380_fragment |  |  |  |  |  |
| PaWa137m_CH35J_006576 \|<br>CH35J_006576_Colletotrichum_<br>higginsianum_fragment | Patatin_and_cPLA2<br>superfamily |  |  |  | Phospholipase A2 |
| PaWa137m.02404 \|<br>PODCO_502086_Fragment | DDE_Tnp_ISL3<br>superfamily |  |  |  | PKS-NRPS hybrid synthetase |
| PaWa137m.02405 \| Pa_4_1058_homolog |  |  |  |  |  |
| gene00315 \| Spok4 |  |  |  |  | Spore killer |
| PaWa137m.Pa_5_2010_fragment1 \|<br>Pa_5_2010_fragment1 |  |  |  |  | GPI anchored serine-threonine<br>rich protein |
| gene01852 \| Uhera2 |  |  |  |  |  |
| PaWa137m_Pa_1_14730_fragment1 \|<br>Pa_1_14730_fragment |  |  |  |  | EST/SMG-like protein 1 |
| gene01815\|Pa_6_9560_Fragment | p450 superfamily |  |  |  | pisatin demethylase |
| gene01816\|Pa_6_9560_homolog | p450 superfamily |  |  |  | pisatin demethylase |
| PaWa137m_7_6640_fragment\|<br>Pa_7_6640_Fragment | Peptidase_M16_M<br>superfamily |  |  |  | insulysin |
| PaWa137m_CTA1_5148\|<br>PaWa137m_CTA1_5148_<br>Colletotrichum_tanacetii_fragment | NK superfamily |  |  |  | Uridylate kinase |
| PaWa137m.gene.002245\|<br>Pa_6_9580_homolog |  |  |  |  | Satratoxin biosynthesis SC1<br>cluster protein 4 |
| gene01818\|Pa_4_3510_homolog | Peptidase_C48<br>superfamily | GumC<br>superfamily |  |  |  |
| PaWa137m.gene.002249\|<br>Pa_1_13070_fragment | zf-H2C2_2 | zf-C2H2 | COG5048 |  | cutinase G-box binding protein |
| PaWa137m.gene.PODCO_102140\|<br>PODCO_102140_fragment | ABC_ATPase<br>superfamily |  |  |  | Heavy metal tolerance protein |
| gene01820\|PV05_03319_<br>Exophiala_xenobiotica_fragment | MdIB |  |  |  | Heavy metal tolerance protein |
| PaWa137m.02410\| Pa_6_7965_homolog |  |  |  |  |  |
| gene01823\|Pa_1_13070_homolog | zf-C2H2 | zf-H2C2_2 | zf-C2H2 | lambda-1<br>superfamily | cutinase G-box binding protein |
| PaWa137m_Pa_6_2490_fragment\|<br>Pa_6_2490_fragment |  |  |  |  |  |
| gene01824\| PODCO_511675_homolog |  |  |  |  |  |
| PaWa137m_Pa_6_2440_fragment\|<br>Pa_6_2440_fragment | PRK09605<br>superfamily |  |  |  |  |
| PaWa137m_MBR_09427_<br>Metarhizium_brunneum_fragment\|<br>PaWa137m_MBR_09427_<br>Metarhizium_brunneum_fragment |  |  |  |  | mitofusin |
| gene01824.2\|<br>PODCO_511675_homolog_2 |  |  |  |  |  |
| PaWa137m_Pa_1_14730_fragment |  |  |  |  | EST/SMG-like protein 1 |
| PaWa137m.02414\|S40285_10826_<br>Stachybotrys_chlorohalonata_homolog |  |  |  |  |  |
| PaWa137m.02415\|<br>Micbo1qcDRAFT_170078_<br>Microdochium_bolleyi_homolog |  |  |  |  |  |
| PaWa137m.02416\| Pa_5_2010_homolog |  |  |  |  | GPI anchored serine-threonine<br>rich protein |
| PaWa137m.gene.002259\|<br>Pa_6_8910_homolog | AdoMet_MTases<br>superfamily |  |  |  | Trans-aconitate 2-methyltransferase |
| gene01828\|CHGG_04275_<br>Chaetomium_globosum |  |  |  |  |  |
| gene01830\|DL98DRAFT_554023_<br>Cadophora_sp_homolog |  |  |  |  |  |
| gene01831\|MBR_09427_<br>Metarhizium_brunneum_homolog |  |  |  |  | mitofusin |
| WaPa137m_4_1930\|<br>Pa_4_1930_fragment | Isoprenoid_Biosyn<br>_C1 superfamily |  |  |  | Iridoid synthase |
| PaWa137m.gene.002264\| CHGG_09441_<br>Chaetomium_globosum_homolog |  |  |  |  |  |
| PaWa137m_NA56DRAFT_749014_<br>Pezoloma_ericae_fragment\|<br>PaWa137m_NA56DRAFT_749014_<br>Pezoloma_ericae_fragment |  |  |  |  |  |
| PaWa137m.gene.002265\|PV05_03319_<br>Exophiala_xenobiotica_fragment | ABC_ATPase<br>superfamily |  |  |  | ABC transporter |
| gene01834\|SPI_07993_homolog_<br>Metallophosphoesterase | MPP_Dcr2 |  |  |  | Metallo-dependent<br>phosphatase |
| PaWa137m_K432DRAFT_223816_<br>Lepidopterella_palustris_homolog\|<br>PaWa137m_K432DRAFT_223816_<br>Lepidopterella_palustris_homolog | TPR_12 |  |  |  |  |
| gene01835\|Pa_1_1830_homolog | GT2_HAS |  |  |  | Hyaluronan synthase |
| gene01836\|Pa_1_14730_homolog | EST1_DNA_bind<br>superfamily |  |  |  | EST/SMG-like protein 1 |
| PaWa137m.02425\|S7711_10142_<br>Stachybotrys_chartarum_homolog | CD_CSD<br>superfamily |  |  |  |  |
| PaWa137m_HIM_12183\|HIM_12183_<br>Hirsutella_minnesotensis_homolog | Pkinase<br>superfamily |  |  |  |  |
| PaWa137m_M441DRAFT_90628\|<br>M441DRAFT_90628_<br>Trichoderma_asperellum_homolog |  |  |  |  |  |
| gene01838\|S7711_11475_<br>Stachybotrys_chartarum_homolog |  |  |  |  |  |
| gene01839\|S40293_11413_<br>Stachybotrys_chartarum |  |  |  |  | patatin-like serine hydrolase |
| gene01840\|NA56DRAFT_703776_<br>Pezoloma_ericae_homolog | DUF3723<br>superfamily | PTZ00121<br>superfamily |  |  |  |
| gene01841\|Micbo1qcDRAFT_129033_<br>Microdochium_bolleyi_homolog |  |  |  |  | Satratoxin biosynthesis SC1<br>cluster protein 4 |
| gene01842\|Pa_5_3190_homolog | MFS superfamily |  |  |  | Efflux pump FUS6 |
| gene01843\|Pa_7_6640_homolog | Ptr superfamily |  |  |  | insulysin |
| PaWa137m_SS1G_08066_Sclerotinia_<br>sclerotiorum_homolog\|Scoty |  |  |  |  |  |
| gene01844\|Chekof | PKc_like<br>superfamily |  |  |  |  |
| PaWa137m_Pa_1_1830_fragment | Peptidase_C48<br>superfamily |  |  |  |  |
| PaWa137m.gene.002278\|PpBr36_10652_<br>Pyricularia_pennisetigena_homolog |  |  |  |  |  |
| gene01845\|Pa_4_3510_fragment |  |  |  |  |  |
| gene01847\|GE09DRAFT_1263105_<br>Coniochaeta_sp_homolog | MdIB |  |  |  | Heavy metal tolerance protein |
| gene01849\|CHGG_09572_<br>Chaetomium_globosum_fragment |  |  |  |  | Putative AC transposase |
| gene01850\|MBR_09427_<br>Metarhizium_brunneum_homolog |  |  |  |  |  |
| PaWa137m_AJ78_06594_<br>Emergomyces_pasteurianus_homolog\|<br>PaWa137m_AJ78_06594_<br>Emergomyces_pasteurianus_homolog |  |  |  |  |  |
| PaWa137m.02432\| Pa_5_3190_homolog | MFS superfamily |  |  |  | Efflux pump FUS6 |
| PaWa137m_CSUB01_08519_<br>Colletotrichum_sublineola_homolog | CFEM superfamily |  |  |  | Satratoxin biosynthesis SC1<br>cluster protein 4 |
| PaWa137m.gene.002286\|<br>MYCTH_64320_<br>Thermothelomyces_thermophilus_<br>homolog |  |  |  |  |  |
| PaWa137m_DL765_007700_<br>Monosporascus_sp_fragment\|<br>PaWa137m_DL765_007700_<br>Monosporascus_sp_fragment |  |  |  |  |  |

